# Improving Accuracy of Somatic Mutation Profiling in Large Epidemiologic Studies: Addressing Cases without Matched Normal Samples

**DOI:** 10.1101/2024.10.28.617052

**Authors:** Arshi Arora, Li Luo, Caroline E. Kostrzewa, Allison Reiner, Venkatraman Seshan, Marc S. Ernstoff, Sharon N. Edmiston, Kathleen Conway, Ivan Gorlov, Klaus Busam, Irene Orlow, Eva Hernando, Nancy E. Thomas, Marianne Berwick, Colin B. Begg, Ronglai Shen, InterMEL

## Abstract

Ideally, detection of somatic mutations in a tumor is accomplished using a patient-matched sample of normal cells as the benchmark. In this way somatic mutations can be distinguished from rare germline mutations. In large retrospective studies, archival tissue collection can pose challenges in obtaining samples of normal DNA. In this article we propose a protocol that improves somatic mutation analysis in the absence of a matched normal sample. The method was motivated by the InterMEL study, a large-scale epidemiologic investigation involving multiomic, multi-institutional genomic profiling of 1000 primary melanoma samples. The key insight for accomplishing improved mutation calling is the fact that germline mutations should produce a variant allele frequency (VAF) of around 50%. While a similar VAF of 50% would also be expected for somatic mutations in pure tumor samples, typically the tumor purity is much less than 50%, resulting in a considerably lower VAF. Making use of a technique that can simultaneously estimate both tumor purity and VAF from tumor-only samples we have developed a method for better distinguishing somatic versus germline variants. Based on 137 melanomas from the InterMEL Study with matched normal tissue to provide a gold standard we show that the conventional pipeline using a panel of (unmatched) normal samples has a false positive rate of 15.6% and a false negative rate of 3.5%. Our new technique improves these error rates to 6.4% and 2.1%, respectively.

## Introduction

Analysis of somatic and germline mutations is increasingly being integrated into routine clinical care for personalized cancer diagnosis and treatment decisions^1-3^. Large-scale epidemiologic investigations are also increasingly focused on collecting and processing tumor samples for characterizing the genomic landscape. Somatic mutations are typically detected by matched tumor-normal sequencing with high accuracy^4^. However, a matched normal control is not always available. This is often the case in retrospective studies or studies that use archival tissue. Tumor-only sequencing in the absence of a matched normal sample is challenging and requires careful methodology. The key problem is that in the absence of a normal DNA sample it is difficult to distinguish somatic variants from germline variants. It is common practice to use population genetics databases to filter out commonly occurring and well-known germline variants, although this approach has limitations for less studied populations, with higher germline false positive rates being reported in non-European individuals due to limited diversity in current population genetics databass^5-7^. Perhaps more importantly, this approach fails to identify rare germline variants. Furthermore, the vast majority of somatic variants observed in tumors are rare variants^8^. The net result is the likelihood of overestimation of somatic tumor mutational burden (TMB) due to the false calling of germline mutations as somatic mutations. TMB has been identified in recent years as an important biomarker for patient survival and treatment response to immunotherapy. It has been shown that the consequence of germline false positives from tumor-only sequencing on TMB quantification can greatly limit the application and further study of this potential biomarker in patient care and treatment outcomes^9,10^.

Several methods have been proposed to identify germline false positives using the observed variant allele frequency of the mutation (VAF)^6,11,12^. The key concept is that tumor samples are typically an impure mixture of tumor and normal cells. Since a germline variant will be represented in both normal and tumor cells, the overall VAF should be in the region of 50% regardless of tumor purity. However, for somatic mutations the VAF will be proportional to tumor purity and thus less than 50% (or much lower for subclonal mutations), offering the opportunity to distinguish true somatic mutations solely on the basis of the VAF. However, this relationship also depends on ploidy and local copy number, and so one needs a method that is also able to adjust for these factors.

In this study, we present Glitter (<u>G</u>erm<u>li</u>ne varian<u>t</u> fil<u>ter</u>), a principled technique for distinguishing somatic and germline variants in the absence of a sample of normal DNA, and illustrate its properties using a sample of cases from the InterMEL consortium. We show that the method has good properties in reducing false positives, giving us confidence that we can accommodate cases without normal DNA in this study without compromising its scientific integrity.

## Methods

### The InterMEL Study

The InterMEL Study is an international effort to assemble tumor and normal specimens from a retrospective cohort of 1000 cases of early stage melanoma diagnosed before the current era of widespread use of immune therapies^13^. To be eligible cases must have been diagnosed between January 1st, 1998 and December 31^st^, 2014 with a cutaneous primary tumor stage IIA-IIID, restaged according to the AJCC 8^th^ Edition. The study, which is still in progress, involves collection of archived tumor and counterpart non-tumor tissue (or germline DNA). Tumor H&Es undergo centralized pathology review. Histopathology-guided co-extraction of nucleic acids from archival tissues is conducted and paired tumor-normal sequencing is performed using the MSK-IMPACT™ assay, a clinically validated and FDA approved hybridization capture-based next-generation sequencing assay developed to guide cancer treatment^1,14^. The assay allows the identification of tumor mutations through targeted deep sequencing by capturing all protein-coding exons and select introns of 505 commonly implicated cancer genes.

### Mutation and copy number calling

Somatic mutation calls from MSK-IMPACT™ were generated using *Roslin*, a research bioinformatics pipeline built to be functionally equivalent to the MSK-IMPACT™ clinical reporting workflow that includes manual filtering/annotation and sign-out by molecular oncologists^15^. Somatic mutations calls were performed by the Bioinformatics Core at Memorial Sloan-Kettering Cancer Center. As a conventional approach for calling somatic mutations in the absence of a matched normal sample a pooled normal was used and post-pipeline processing was performed to filter out potential germline variants using the ExAC database^16^ (ExAC allele frequency >=0.005) and sequencing artifacts using a panel of normal samples selected for enrichment for recurrent artifacts. Copy number analysis was performed using the FACETS algorithm^17^. For matched tumor-normal sample pairs, joint segmentation and integer copy number estimation was accomplished from joint analysis of total copy ratio logR and allelic copy ratio logOR. The logR is computed from the total sequence read count at all SNP sites in the tumor versus normal and logOR computed from the variant-allele count at heterozygous loci in tumor versus the matched normal. For tumor-only copy number analysis, we use an unmatched normal sample that is similar in sequence coverage to the tumor sample by quantile-matching. The FACETS algorithm permits estimation of the integer copy number and the tumor sample purity, parameters that are crucial for our method for calling somatic mutations in cases without a matched normal sample (see next section).

### Method for identifying false positive somatic variants

Let *v* denote the observed variant allele frequency at a given nucleotide. This is considered to be a binomial random variable among the total of *n* reads at a given nucleotide, *Bi*(*n*, θ). For a germ-line heterozygous variant θ = 1/ 2, that is the variant should be present in approximately 50% of the *n* reads. In a completely pure tumor sample with normal copy number the VAF has exactly the same distribution, and so there is no leverage for distinguishing a somatic mutation from a germline variant. However, tumors are typically an impure mixture of tumor and normal cells. If the proportion of tumor cells, the sample purity, is denoted by *p*, then θ = *p* / 2.As a result the VAF is expected to be lower than 50% to an extent that is proportional to *p*. Our method involves calling observed variants false positives (i.e. germline variants) if the observed VAF exceeds its upper 95% confidence interval for a somatic variant based on the estimated tumor purity. The preceding simple setting applies only if there is a normal allele copy number of 2. Estimates of copy number and tumor purity are products of the FACETS algorithm. In this more general setting the expected VAF is θ = *pc* / (*pt* + 2(1− *p*)) for a somatic variant, while it is θ = (*pc* + (1− *p*)) / (*pt* + 2(1− *p*)) for a germline variant, where *c* is the variant specific copy number and *t* is the total copy number. In this setting we once again designate the variant a false positive if the VAF falls within the corresponding lower 95% confidence limit for a germline variant in the presence of copy number alterations.

## Results

### Validation study using tumors with normal tissue samples

We examined the accuracy of the method in 137 cases from InterMEL for which matched normal DNA was available. These samples underwent MSK-IMPACT sequencing with a mean coverage of 293 and 239 in tumor and normal DNA respectively. Our gold standard benchmark involved the standard procedure, whereby the matched normal sample was used to filter out germ-line variants. The matched analysis called 5,778 somatic mutations in total with a median of 15 per tumor. Use of the conventional approach for tumor-only samples involving a pooled normal sample led to 6604 mutations with a median of 21 per tumor, representing a significant upward shift in TMB (Figure 1). The false positive rate was observed to be high, at 15.6% (1031/6604), and the false negative rate was 3.5% (205/5778).

**Figure 1:**
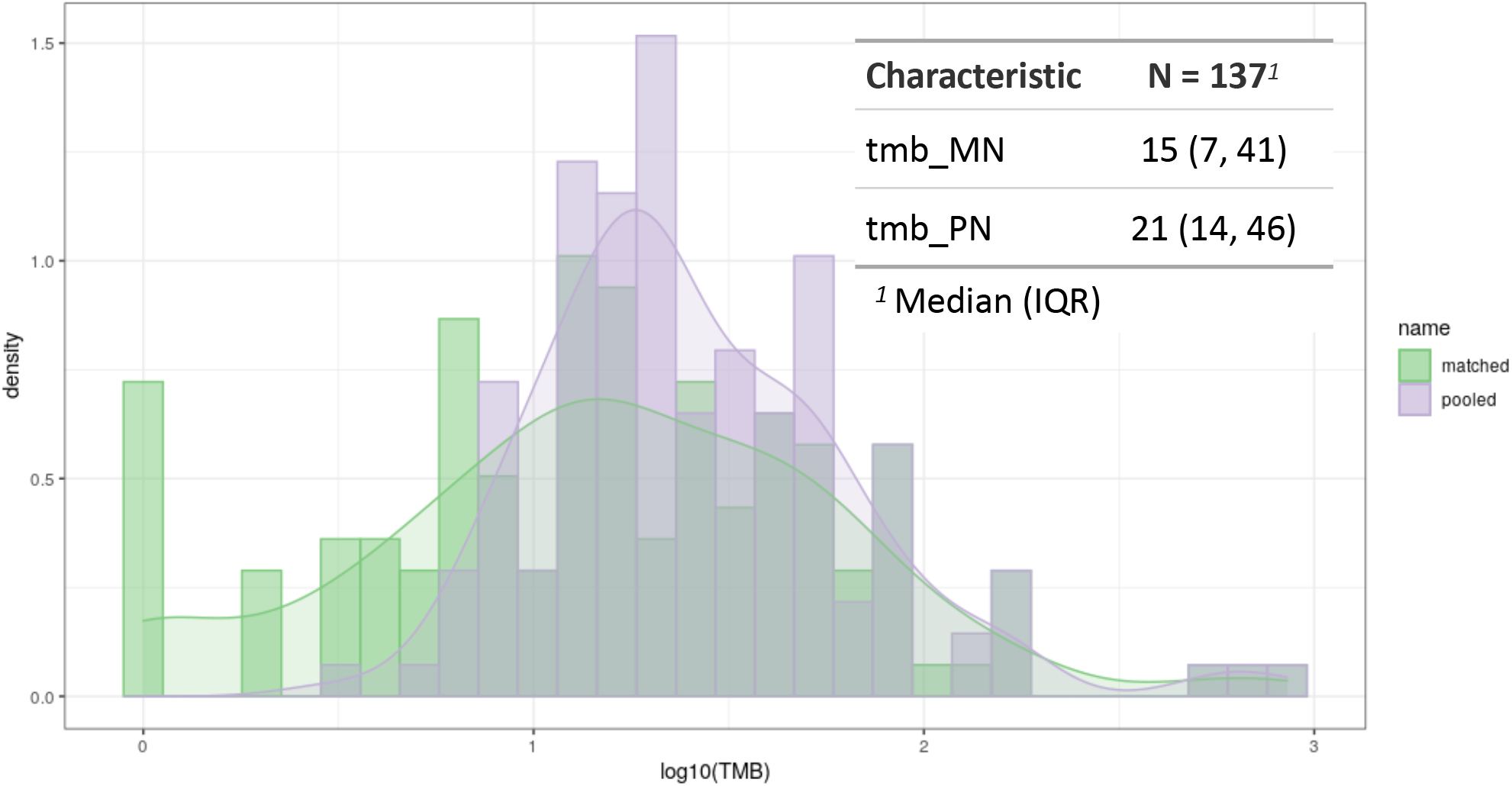
Distribution of log10(TMB) of 137 samples as seen across a typical mutation calling pipeline with matched normal(green) vs a tumor-only pipeline (purple)

Figure 2a shows the distribution of variant allele frequency (VAF). The VAF for false positives has a much higher mean of 0.46 contrasting a mean VAF of 0.2 for true positives. Such difference in VAF diminishes in higher purity tumor samples (Figure 2b). This suggests that a main source of false positive calls in tumor-only mutation analysis may come from rare germline variants insufficiently filtered out by the population database used as a reference.

**Figure 2:**
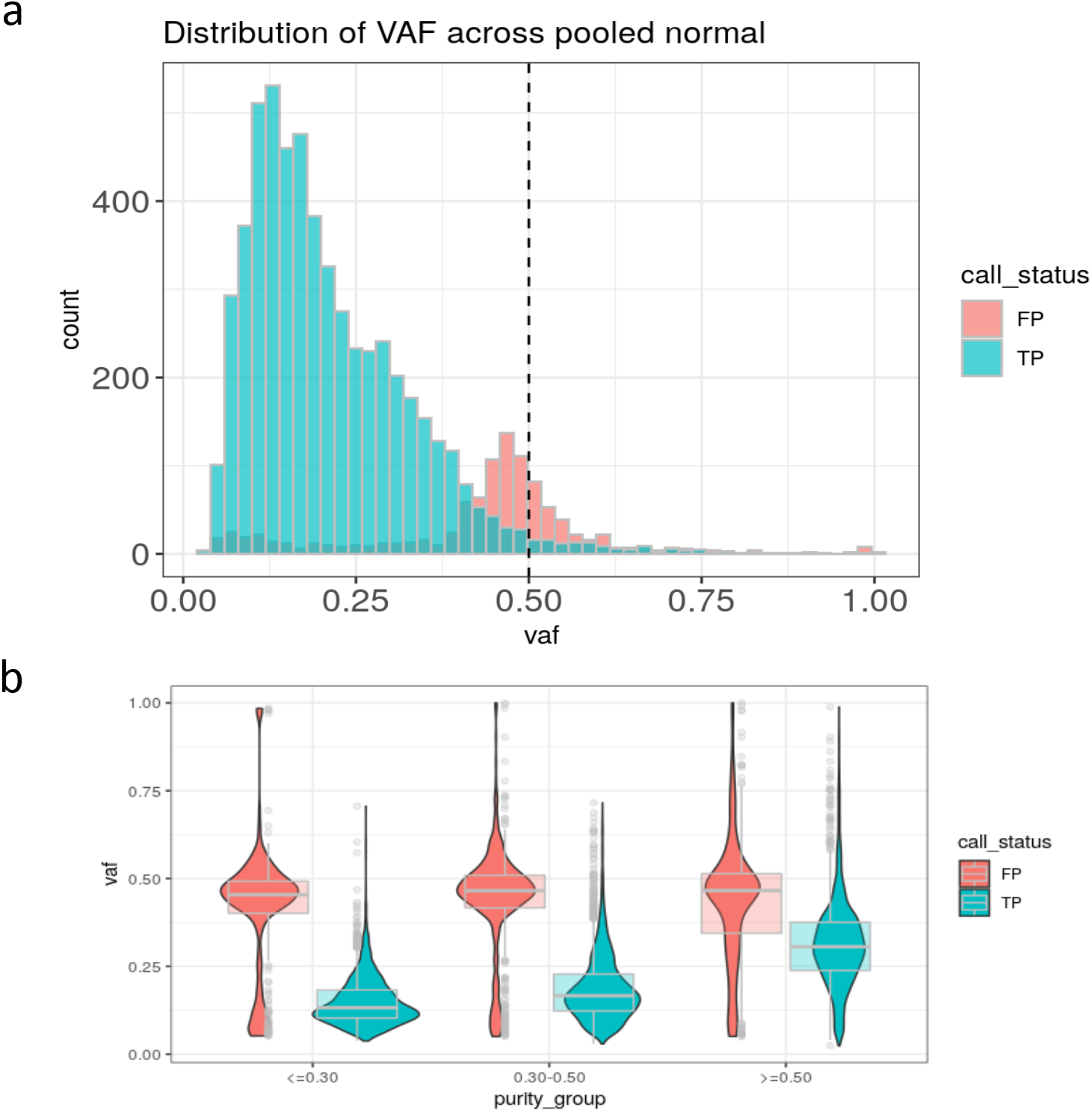
Variant allele frequency distribution stratified by a) false positive (FP) and true positive (TP) and b) further by tumor purity.

When we applied Glitter for filtering out sample-specific germline variants the false positive rate improved to 6.4%, while the false negative rate was 2.1%. These correspond to a sensitivity of 97.8% and a positive predictive value of 92.8%. Table 1 further demonstrates that the performance improves as the tumor purity declines, as we would expect.

**Table 1:**
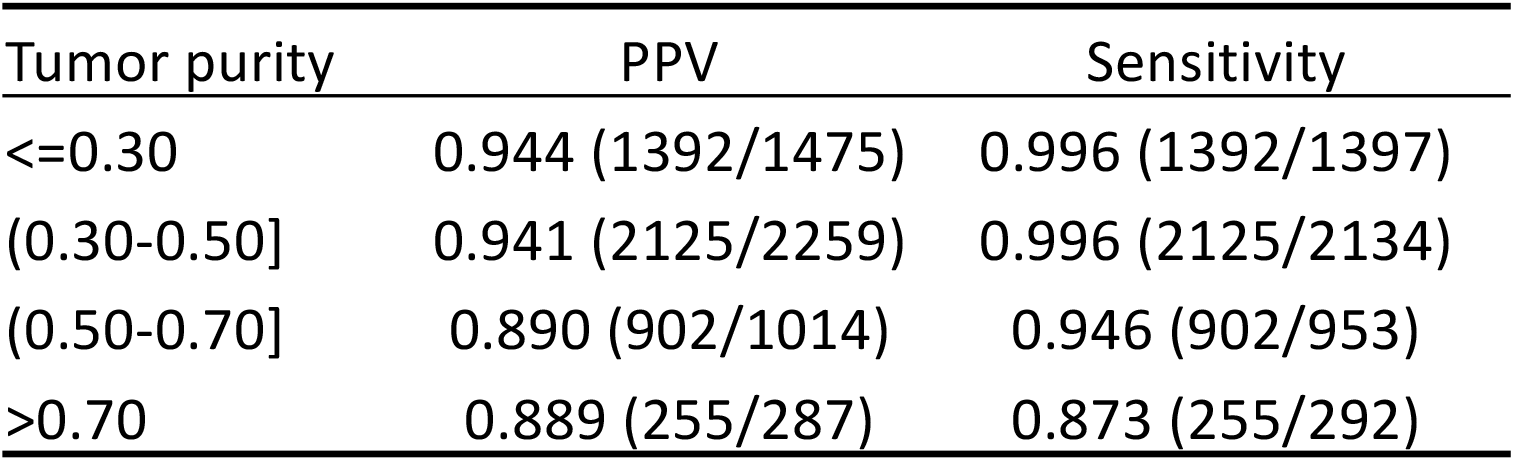
Sensitivity (Recall) and PPV (Precision) for tumor-only mutation analysis with Glitter filtering for different tumor purity. Numerators and denominators are included in parenthesis.

### Application of tumor-only sequencing to cases with no normal tissue

We applied the tumor-only method to the 94 cases assembled to date in InterMEL without matched normal samples. Clearly for these cases we have no benchmark against which to gauge our results. However, we can evaluate the overall called mutational burden to see if it is generally concordant with that obtained in the 137 cases evaluated conventionally with matched normal samples. The median VAF for variants called in these samples is 0.28. There is a clear second mode centered around 0.5 largely representing germline false positive calls (Figure 3a). The distribution of TMBs for unfiltered calls in these 94 tumor-only cases is notably higher than for the 137 cases with normal samples but the discrepancy is largely eliminated after the presumed germline calls are filtered out by our algorithm (Figure 3b).

**Figure 3:**
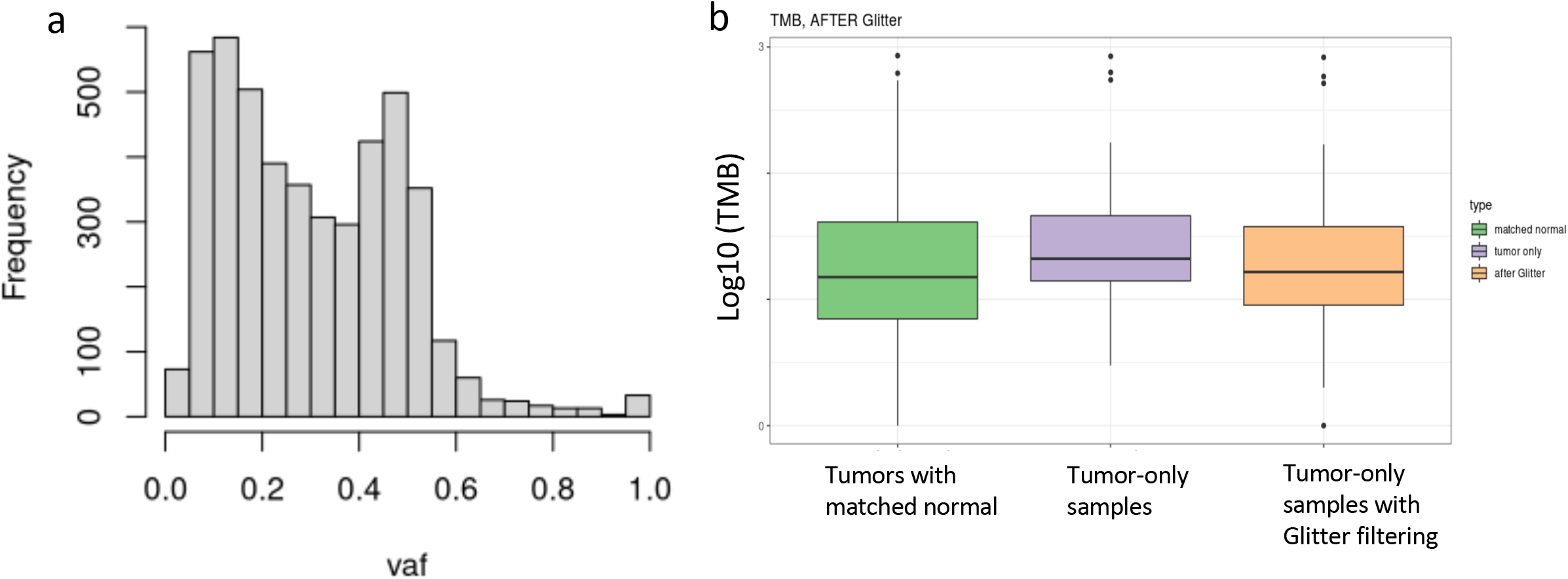
Analysis of 94 tumor-only samples showing a) VAF distribution with a median VAF of 0.28 and a second peak at 0.45; b) log10(TMB) for tumors with matched normal, the tumor-only samples and tumor-only samples with Glitter filtering for germline false positives.

## Discussion

Increasingly, large cancer epidemiologic investigations encompass the collection of tumor tissue samples. These are useful for multiple purposes, including investigation of tumor subtypes defined by somatic profiles, and the exploration of relationships between genetic host factors and somatic features of the tumors. Ideally, such studies should involve collection of samples of matched normal tissue, since these are necessary to definitively distinguish true somatic mutations from private germline variants in the patient. While pooled normal samples can play a useful role for identifying common germline variants, these are of essentially no value for interpreting the vast majority of somatic mutations that are rare variants. In this article we demonstrated how false positive somatic calls due to the presence of a rare germline variant can be filtered out with much greater accuracy.

The work was motivated by the InterMEL Study, a cohort study designed to investigate the natural history of early-stage melanoma. This study involves multi-omic profiling of primary tumor samples of melanoma patients diagnosed with stages IIA-IIID disease. For somatic mutation profiling we used the MSK-IMPACT™ targeted panel, consisting of approximately 500 cancer related genes. The method could easily be applied to different genomic panels. The method does however rely on the use of methods to estimate copy number at the location of each suspected mutation and tumor purity. The InterMEL study is in progress, and its design and early results are described in more detail in Luo et al.^13^. The challenges of conducting multiomic profiling in this large epidemiologic study of tumors that are typically very small with limited DNA are also described in detail in Orlow et al.^18^

## Acknowledgement

This work was supported by the National Cancer Institute at the National Institutes of Health (Grant numbers P01CA206980, R01CA251339), Memorial Sloan Kettering Cancer Center (Grant number P30 CA008748).

## Declaration of Interest

Arshi Arora

Employment: Volastra, New York, NY

Caroline E. Kostrzewa

Stock and Other Ownership Interests: Johnson & Johnson/Janssen

Marc S. Ernstoff

Stock and Other Ownership Interests: Johnson & Johnson/Janssen, Becton Dickinson, Abbott Laboratories, AbbVie, AstraZeneca, Bristol Myers Squibb/ Roche, Lilly, Merck, Thermo Fisher Scientific, United Health Group

Li Luo

Employment: Baxter

Stock and Other Ownership Interests: Baxter

Irene Orlow

Leadership: Tolstoy Foundation Rehabilitation and Nursing Center, Inc No other potential conflicts of interest were reported.

Klaus J. Busam

Patents, Royalties, Other Intellectual Property: Royalties for textbook published by Elsevier

## Key points

- Somatic mutation analysis from tumor-only sequencing can be confounded by rare germline variants, leading to inflated tumor mutational burden estimates.
- In this article we propose a protocol that improves somatic mutation analysis in the absence of a matched normal. The method was motivated by the international InterMEL consortium study, a large-scale epidemiologic study of primary melanoma involving multi-omic, multi-institutional genomic profiling.
- Making use of variant allele frequency (VAF), tumor purity and copy number alterations we have developed a method for better distinguishing somatic versus germline variants.
- Using samples from the InterMEL study with matched normal tissue to provide a gold standard we show that our approach significantly improves upon the conventional pipeline using a panel of (unmatched) normal samples in reducing the error rates.

